# CSF biomarkers are differentially related to structural and functional changes in dementia of the Alzheimer’s type

**DOI:** 10.1101/250449

**Authors:** Charles B Malpas, Michael M Saling, Dennis Velakoulis, Patricia Desmond, Rodney J Hicks, Henrik Zetterberg, Kaj Blennow, Terence J. O’Brien

**Affiliations:** Department of Medicine, Royal Melbourne Hospital, VIC, Australia; Melbourne School of Psychological Sciences, The University of Melbourne, VIC, Australia; Developmental Imaging, Murdoch Children’s Research Institute, Melbourne, VIC, Australia; Department of Psychiatry, University of Melbourne, VIC, Australia; Department of Radiology, University of Melbourne, VIC, Australia; Centre for Molecular Imaging, Peter MacCallum Cancer Centre, Melbourne, VIC, Australia; Department of Psychiatry and Neurochemistry, the Sahlgrenska Academy at the University of Gothenburg, Mölndal, Sweden; Clinical Neurochemistry Laboratory, Sahlgrenska University Hospital, Mölndal, Sweden; Department of Molecular Neuroscience, UCL Institute of Neurology, Queen Square, London, United Kingdom; UK Dementia Research Institute at UCL, London, United Kingdom

**Keywords:** Alzheimer’s, P-tau, beta-amyloid, cerebrospinal fluid, cortical thickness, cerebral glucose uptake

## Abstract

The two cardinal pathologies of Alzheimer’s disease (AD) develop according to distinct anatomical trajectories. Cerebral tau-related pathology first accumulates in the mesial temporal region, while amyloid-related pathology first appears in neocortex. The eventual distributions of these pathologies reflect their anatomical origins. An implication is that the cardinal pathologies might exert preferential effects on the structurofunctional brain changes observed in AD. We investigated this hypothesis in 39 patients with dementia of the Alzheimer’s type. Interrelationships were analysed between cerebrospinal fluid (CSF) biomarkers of the cardinal pathologies, volumetric brain changes using magnetic resonance imaging, and brain metabolism using [^18^F]-FDG-PET. Amyloid-related pathology was preferentially associated with structurofunctional changes in the precuneus and lateral temporal regions. Tau-related pathology was not associated with changes in these regions. These findings support the hypothesis that tau- and amyloid-pathology exert differential effects on structurofunctional changes in the AD brain. These findings have implications for future therapeutic trials and hint at a more complex relationship between the cardinal pathologies and disruption of brain networks.

## Introduction

Alzheimer’s disease (AD) is characterised by two cardinal pathologies, namely the intracellular accumulation of tau-related neurofibrillary tangles (NFTs), and the accumulation of various soluble and insoluble extracellular amyloid-beta (Aβ) aggregates. In typical AD, the genesis of these pathologies follows well-described anatomical trajectories. The formation of cerebral tau-related pathology begins in the entorhinal cortex, then spreads to nearby limbic regions (including the hippocampus) before infiltrating broad regions of isocortex [1]. In contrast, the formation of amyloid-related pathology begins in basal isocortex before spreading inwards to mesial temporal structures and finally involving more diffuse isocortical regions [2]. Several *in vivo* imaging studies have revealed greatest Aβ binding in anterior neocortex, with relatively little uptake in mesial temporal structures [3-5].

The different anatomical distribution of tau- and amyloid-pathology raise the question of whether they preferentially relate to structurofunctional changes in the AD brain. We have previously hypothesised that tau-pathology is preferentially associated with changes in the mesial temporal regions, while amyloid pathology is preferentially associated with diffuse neocortical changes [6, 7]. The aim of this study was to investigate this hypothesis using volumetric measurements obtained via structural magnetic resonance imaging and relative cerebral metabolism assessed with [^18^F]-FDG-PET. These modalities have been demonstrated to be sensitive to the pathological processes of AD [8]. Cerebrospinal fluid (CSF) protein markers of AD pathology were used as measures of the cardinal pathologies, as they present the only widely available method of measuring P-tau and Aβ pathological load *in vivo* using a common modality [9, 10].

Here we report an investigation of this preferential association hypothesis in a cohort of patients with dementia of the Alzheimer’s type (DAT). We expected to find differential relationships between CSF biomarkers and regional structurofunctional changes in AD. Specifically, CSF levels of P-tau were expected to be preferentially associated with structurofunctional changes in the mesial temporal structures. Levels of Aβ in CSF were expected to be preferentially associated with structurofunctional changes in neocortex.

## Material and Methods

### Participants

Participants were 39 patients (22 males, 17 females) with DAT, diagnosed according to National Institute on Aging – Alzheimer’s Association criteria [11]. All were recruited from clinical sources in Melbourne, Australia. Ethics approval was obtained from the institutional ethics committee (Melbourne Health, HREC 2012.148) and all patients gave written informed consent. We have reported resting-state fMRI findings from this cohort elsewhere [7].

### CSF Biomarkers

CSF biomarkers were sampled via lumbar puncture, aliquoted and stored at -80°C pending analyses. CSF Aβ1-42 levels were determined using a sandwich ELISA method (INNOTEST^®^ ß- AMYLOID(1-42), Innogenetics, Gent, Belgium), as previously described [12]. CSF total tau (T-tau) concentrations was determined using a sandwich ELISA (Innotest hTAU-Ag, Innogenetics, Gent, Belgium) specifically constructed to measure all tau isoforms irrespectively of phosphorylation status [13], while CSF P-tau (phosphorylated at threonine 181) was measured using a sandwich ELISA method (INNOTEST^®^ PHOSPHO-TAU(181P), Innogenetics, Ghent, Belgium), as described previously in detail [14]. Cut-offs for the analyses were 530 pg/mL for Aβ1-42, 350 pg/mL for T-tau and 60 pg/mL for P-tau [15]. All CSF measurements were performed in one round of experiments using one batch of reagents, by board-certified laboratory technicians, who were blinded to clinical data. Internal quality control (QC) samples were run on each plate to assure consistency. Intra-assay coefficients of variation were < 11.1% for Aβ1-42, <13.3% for T-tau and <9.2% for P-tau.

### Image Acquisition

Structural images were acquired on a 3.0T Siemens Tim Trio scanner at the Royal Melbourne Hospital, Melbourne, Australia. Structural images comprised a 176-slice sagittal 3D acquisition (MPRAGE; flip angle = 9**°**, TR = 1900 ms, TE = 2.13 ms, TI = 900 ms, FOV = 176 × 256, matrix = 256 × 256; slice thickness = 1 mm). All image volumes were inspected at the time of acquisition, with acquisition repeated were necessary (e.g., due to gross movement artefact).

[^18^F]-FDG-PET images were acquired at the PET Centre at the Peter MacCallum Cancer Centre in Melbourne. After a minimum of 6 hours fasting, patients were instructed to lie quietly in a darkened room for 15 mins. Patients were then injected with 220 MBq of fluorodeoxyglucose ([^18^F]-FDG-PET) through an intravenous catheter and continued resting for at least 45 mins. After resting the patients were scanned on a GE Discovery 690 (GE Medical Systems Milwaukee, WI). Patients were positioned supine with their heads secured in a dedicated head-rest. A low dose CT scan for attenuation correction was acquired with the following exposure parameters, 120kV, Auto mA range 40-80, rotation time of 0.5 sec, pitch 0.984, slice thickness 3.75 mm. A 15-minute list-mode PET acquisition of the brain was then acquired and re-framed into a dynamic scan of 15 × 1 minute frames to detect any patient motion. Frames with motion present were excluded from processing. All PET scans were processed without time-of-flight using OSEM3D iterative reconstruction using 8 iterations, 24 subsets, 5 mm filter, 192 matrix and a 35 cm field of view. Images were inspected following acquisition to ensure no gross artifact (i.e., movement) rendered them unusable.

### Image Processing

Cortical thickness analysis was using the Freesurfer software package (version: 5.2.0), the full technical details of which are described elsewhere [16]. The resulting cortical models were visually inspected by an experience operator (CBM) to ensure accuracy. Manual corrections to the brain mask were made where non-cortical tissue had been included in the cortical model. Six regions of interest were specified in order to extract mean thickness estimates from the cortical model. These included (a) pre-frontal cortex, (b) orbitofrontal cortex, (c) precuneus, (d) lateral-temporal cortex, (e) anterior-cingulate cortex, and (f) entorhinal cortex. These regions have been previously identified as containing high amyloid-load using in vivo PET imaging [17]. The anterior cingulate was included following previous work from our group implicating this region in amyloid-beta load [7]. The specific ROIs (and their FreeSurfer labels) for each region are shown in *Figure 1*. Cortical thickness estimates were averaged across ROIs and also across hemispheres, resulting in a single mean thickness estimate for each region. For the hippocampus, the estimated volume was extracted from the FreeSurfer sub-cortical pipeline and averaged across hemispheres.

**Figure 1.**
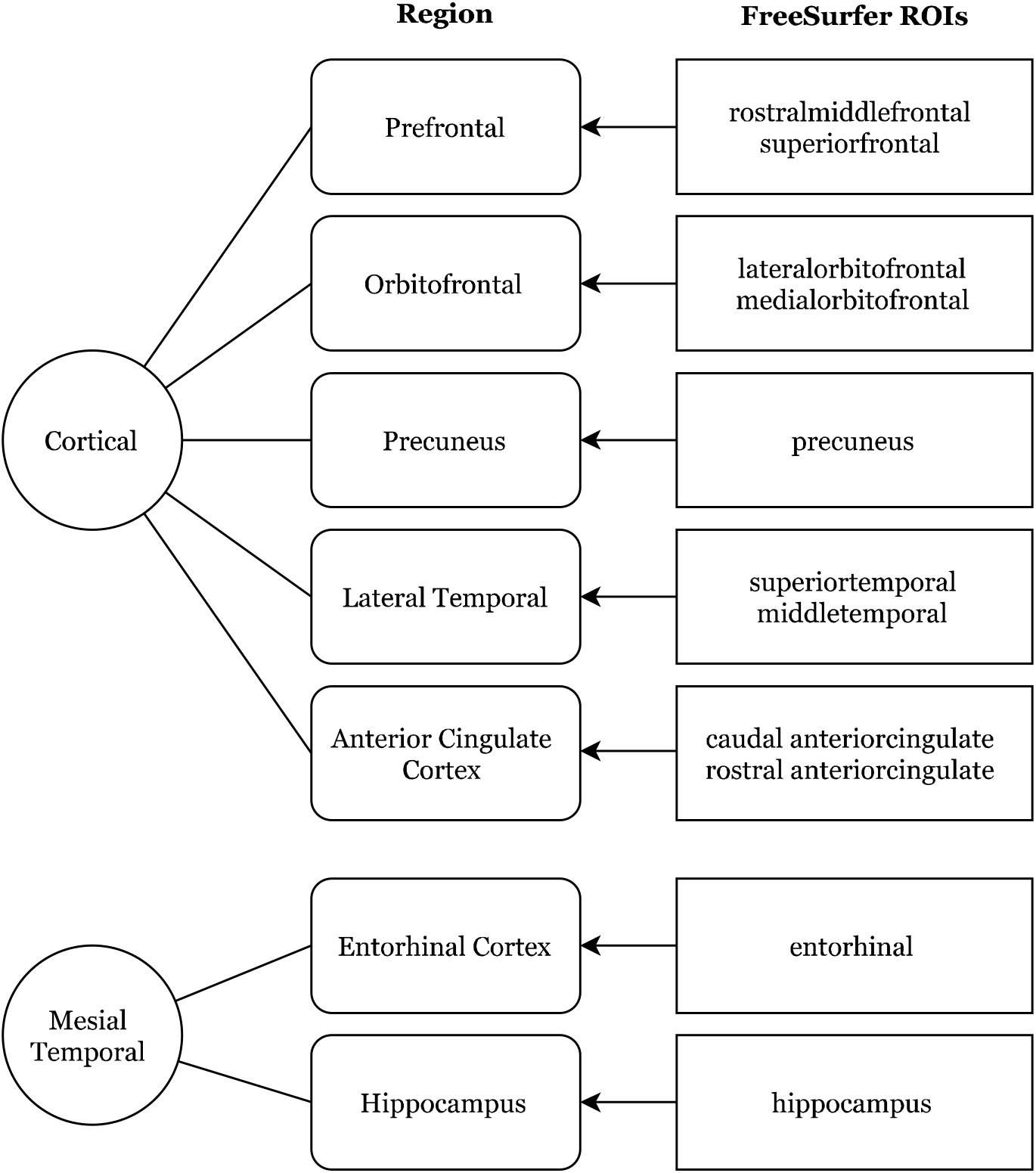
Specific FreeSurfer regions of interest. The FreeSurfer ROIs are given as the standard label names used in the software.

The [^18^F]-FDG-PET images were aligned to FreeSurfer structural space using a linear boundary-based registration approach [18]. Briefly, this involves segmenting the structural reference image into tissue classes and coregistering the FDG-PET image by maximizing the intensity gradient across tissue boundaries. The transformation matrices were then used to inversely register the FreeSurfer ROIs to native [^18^F]-FDG-PET space. In this way, the [^18^F]-FDG-PET volumes were not subjected to potential degradation associated with manipulation or interpolation. Mean [^18^F]-FDG-PET uptake was extracted from each of the six cortical regions and the hippocampus as described above. In addition, the mean uptake for the entire cerebellum was extracted. Normalised mean uptake for each ROI was then calculated by dividing by the mean uptake for the cerebellum.

### Statistical Analysis

All analyses were performed in SPSS v24.0.0 (IBM Corporation). The relationship between CSF biomarkers and imaging metrics was analysed using Pearson’s partial correlation coefficients. For [^18^F]-FDG-PET, mean uptake in each of the seven regions was correlated with each of the three biomarkers, adjusted for age. For the structural analyses, mean cortical thickness for the six pial regions, and the volume of the hippocampus, were correlated with the three biomarkers adjusting for age and intracranial volume (ICV). ICV was computed using the default procedure implemented in FreeSurfer. Briefly, this procedure estimates ICV by exploiting the known relationship between true intracranial volumes and the determinant of transform matrix produced by normalizing the whole-brain T1 image to MNI305 space. As described in Buckner and colleagues [19], this approach has been validated against manually segmented intracranial volume and is minimally biased in populations with dementia.

This resulted in 42 statistical comparisons. Given the increased risk of type 1 error, the Benjamini-Hochberg procedure was used to control the false discovery rate (FDR) at 5% [20]. FDR correct p-values below .05 were considered statistically significant. In order to show a true statistical dissociation (i.e., an interaction), it is necessary to demonstrate that the two correlation coefficients are statistically significantly different from one another, not merely that one is statistically significantly different from zero and the other is not [21]. In order to achieve this, where a statistically significant region was found the correlation between it and Aβ was compared to the correlation with P-tau using an approach described elsewhere for dependent correlation coefficients [22]. Statistically significant correlations were subjected to sensitivity analysis to determine the effect of Aβ status. Specifically, the sample was split into those with ‘normal’ versus ‘abnormal’ CSF Aβ1-42 concentrations (defined as Aβ1-42 < 530 pg/mL, as described above). Correlation coefficients were then computed for the ‘abnormal’ sub-sample in isolation.

## Results

### Descriptive Statistics

Descriptive data for the study sample are shown in *Table 1*.

**Table 1.** Sample characteristics

|  | *Mean* | *95% CI* | *Range* |
| --- | --- | --- | --- |
| *Demographics* |  |  |  |
| Age (Years) | 70 | 68 - 73 | 57 - 83 |
| MMSE (Total) | 20 | 19 - 21 | 14 - 29 |
| ADAS-Cog (Total) | 20 | 17 - 22 | 7 - 31 |
| *CSF Biomarkers* |  |  |  |
| P-tau (pg/ml) | 86 | 75 - 96 | 27-175 |
| T-tau (pg/ml) | 940 | 806 - 1075 | 192 - 1962 |
| Aβ1-42 (pg/ml) | 481 | 413 - 549 | 224 - 1332 |

*N* = 39 (22 males, 17 females). MMSE = Mini Mental Status Examination, ADAS-Cog =
Alzheimer's Disease Assessment Scale – Cognitive

**Table 2.** Descriptive data for the study sample

|  | *Mean [95% CI]* |  |
| --- | --- | --- |
|  | *Structural volumetrics* | *FDG-PET uptake* |
| Prefrontal | 2.38 [2.33, 2.42] | 1.14 [1.11, 1.17] |
| Orbitofrontal | 2.36 [ 2.31, 2.42] | 1.03 [1.00, 1.05] |
| Precuneus | 2.03 [1.97, 2.10] | 1.12 [1.07, 1.17] |
| Lateral Temporal | 2.47 [2.41, 2.52] | 0.93 [0.90, 0.96] |
| Anterior Cingulate<br>Cortex | 2.72 [2.65, 2.80] | 0.97 [0.95, 0.99] |
| Entorhinal Cortex | 2.77 [2.63, 2.90] | 0.68 [0.65, 0.70] |
| Hippocampus | 3063.78 [2866.78,<br>3261.37] | 0.81 [0.79, 0.84] |

*Note:* Structural volumetrics are presented in millimetres (mm) for all measures, except for the hippocampus which is presented in millimetres cubed (mm<sup>3</sup>). FDG-PET uptake values are normalised to the cerebellum.

### Cortical Thickness

As shown in *Figure 2*, mean cortical thickness in the precuneus was positively associated with CSF levels of Aβ1-42, *r* = .49, *p* = .01. There was no statistically significant relationship with P-tau, *r* = -.14, *p* = .91. The difference between these correlation coefficients was statistically significant, *t*(36) = 2.97, *p* = .005. A similar pattern was observed in the lateral temporal region. Mean cortical thickness was positively correlated with CSF Aβ1-42 concentration, *r* = .49, *p* = .01, but there was no statistically significant correlation with P-tau, *r* = -.11, *p* = .86. The difference between these correlation coefficients was statistical significance, *t*(36) = 2.82, *p* = .008. As shown in *Figure 4*, no statistically significant associations emerged for T-tau.

**Figure 2.**
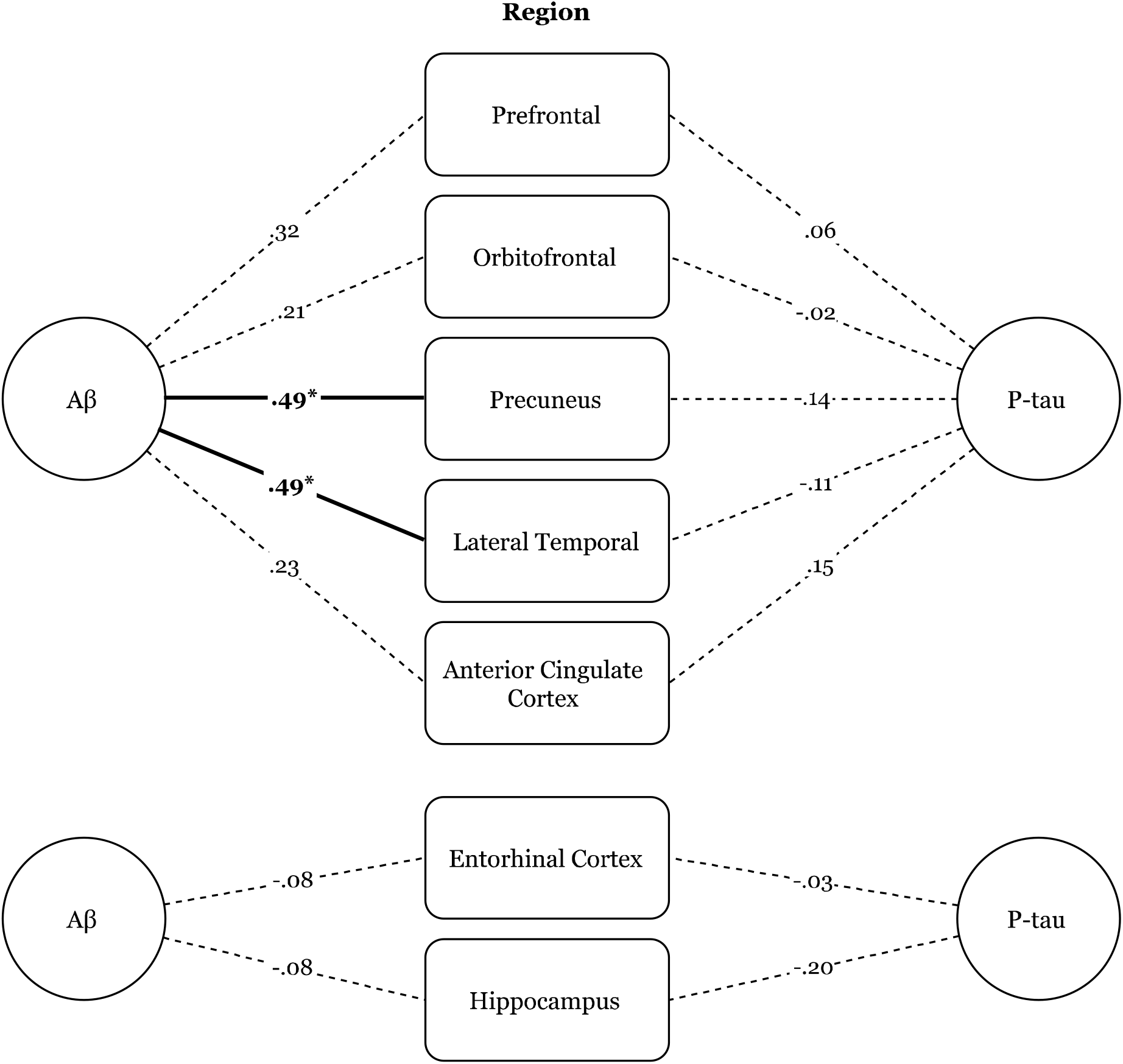
Partial correlation between CSF biomarkers and structural volumetrics in specific regions. Only the precuneus (*p* = .01) and lateral temporal (*p* = .01) regions were correlated with Aβ, while no regions were correlated with P-tau. Asterisk indicate correlations that are statistically significant at *p <* .05 (FDR corrected).

### [_18_F]-FDG-PET

Only two regions showed statistically significant correlations with the CSF biomarkers. These results are shown in *Figure 3*. Uptake in the precuneus was positively correlated with levels of Aβ1-42, *r* = .44, *p* = .02. The correlation with P-tau did not reach significance, *r* = -.07, *p* = .97. The difference between these correlation coefficients was statistically significant, *t*(36) = 2.32, *p* = .03. The correlation between the Aβ1-42 and FDG uptake in the lateral temporal region was also positive and statistically significant, *r* = .44, *p* = .02. The correlation with P-tau and uptake in this region did not reach significance, *r* = -.01, *p* = .97. Again, the difference between these coefficients was statistically significant, *t*(36)= 2.09, *p* = .04. As shown in *Figure 4*, no statistically significant associations emerged for T-tau (all *ps >* .05).

**Figure 3.**
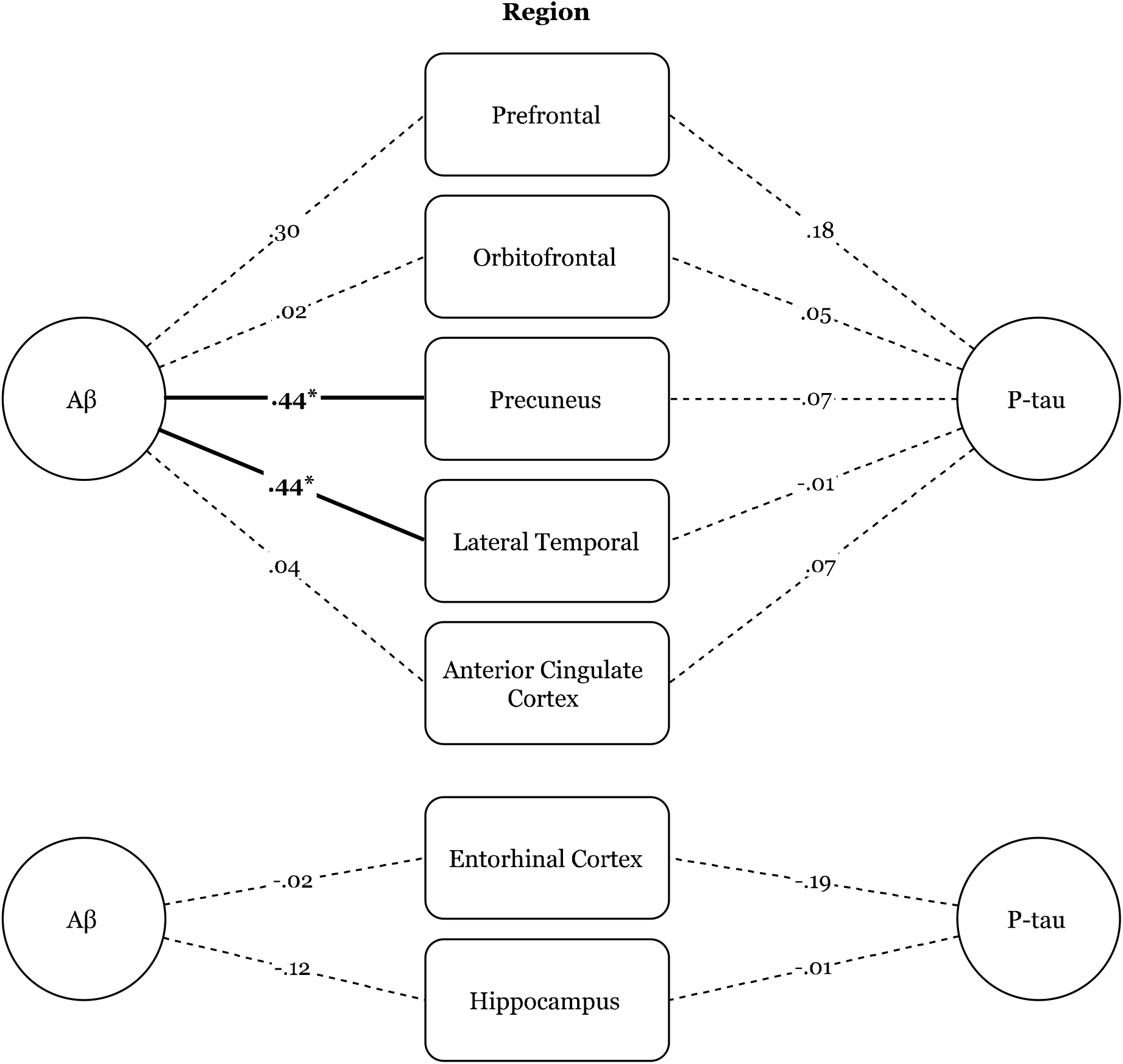
Partial correlation between CSF biomarkers and FDG-PET uptake in specific regions. Only the precuneus (*p* = .02) and lateral temporal regions (*p* = .02) were correlated with Aβ, while no regions were correlated with P-tau. Asterisk indicate correlations that are statistically significant at *p <* .05 (FDR corrected).

**Figure 4.**
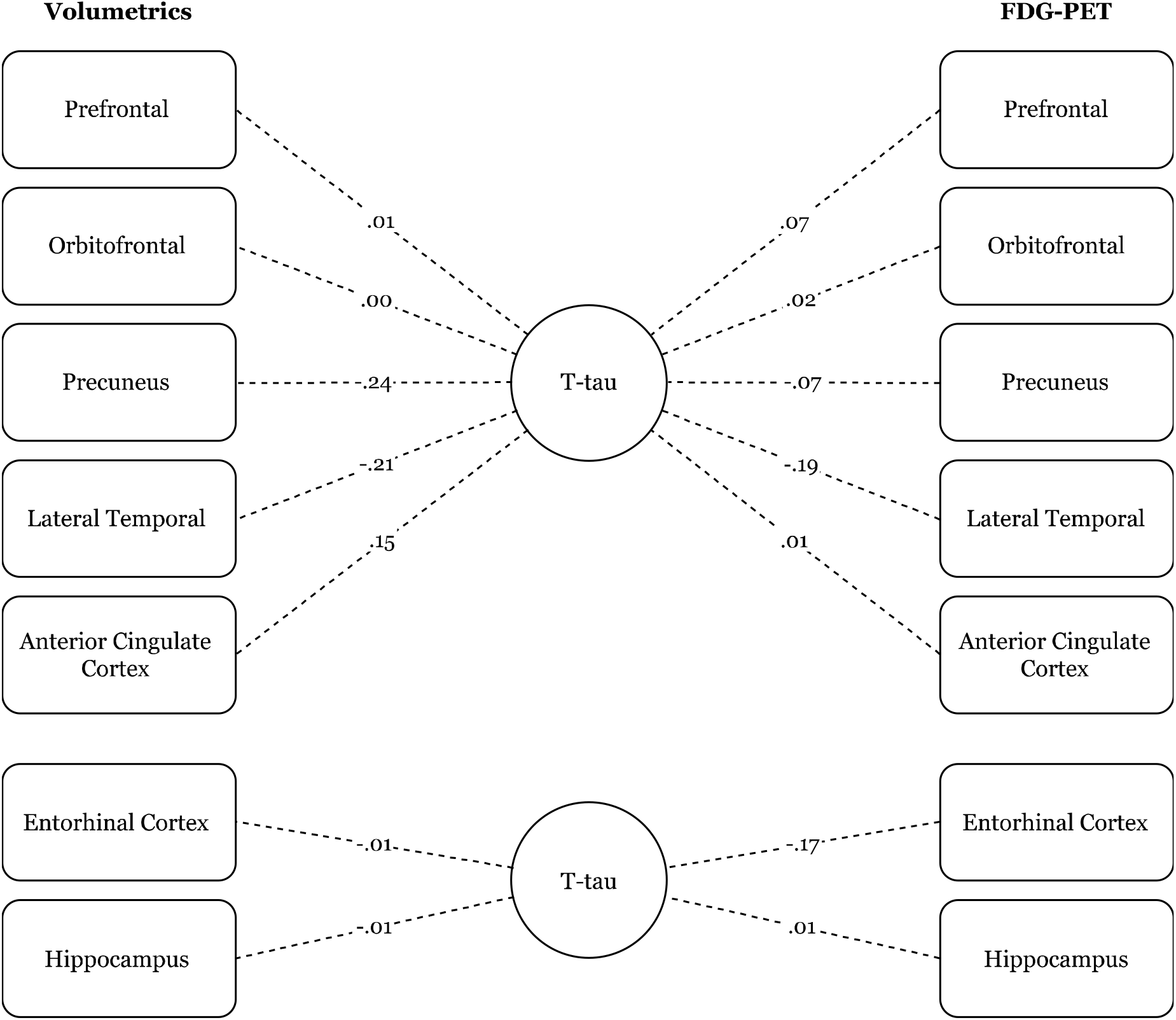
Partial correlation between CSF total tau, FDG-PET uptake, and structural volumetrics in specific regions. No relationships were statistically significant at the *p <* .05 level (FDR corrected).

### Effect of CSF Aβ1-42 status

Twenty-nine participants (74%) had abnormal CSF Aβ1-42 concentrations. Despite the smaller sample size, the correlation between Aβ1-42 and FDG uptake in the precuneus remained statistically significant, *r* = .46, *p* = .013. The correlation between Aβ1-42 and FDG uptake in the lateral temporal regions, however, was no longer statistical significance, *r* = .14, *p* = .48. The correlation between CSF Aβ1-42 and cortical thickness in the precuneus was slightly reduced in the sub-sample and no longer, *r* = .33, *p* = .10. A similar pattern was observed for thickness in the lateral temporal regions, *r* = .21, *p* = .28. No correlations between CSF P-tau and neuroimaging markers in the sub-sample.

## Discussion

The findings of this study partially support a preferential association between AD biomarkers and structurofunctional changes in patients with DAT. As expected, levels of Aβ-related pathology were associated with [^18^F]-FDG-PET uptake and cortical thickness in the precuneus and lateral temporal regions. While reduced glucose metabolism and cortical thickness in these regions was associated with greater levels of Aβ-related pathology, there was no detectable relationship with tau-related pathology. These differential relationships were statistically supported. As the precuneus and lateral temporal regions are amongst those with greatest Aβ accumulation in AD, these preferential associations are consistent with the hypothesis discussed here. No other regions showed detectable inter-relationships between [^18^F]-FDG-PET uptake, cortical thickness, and disease biomarkers.

Our findings are partially consistent with previous reports. For example, Vukovich and colleagues also reported a positive association between CSF Aβ and glucose metabolism in the right temporal region [23]. As with our study, no relationships were found between glucose metabolism and CSF T-tau. Statistical relationships were, however, reported between CSF Aβ, prefrontal, and anterior cingulate hypometabolism. Chiaravalloti and colleagues also investigated the FDG-PET correlates of CSF biomarkers in Alzheimer’s disease [24]. Higher CSF P-tau and T-tau was associated with reduced uptake in frontal and limbic regions. CSF Aβ was associated with wide-spread cortical dysfunction. While the existence of differential relationships between the biomarkers is broadly consistent with our hypothesis, the regions reported are not consistent with the regions revealed by our study.

In terms of structural imaging studies, our findings are again partially consistent with previous reports in the literature. For example, Ossenkoppele and colleagues [25] found a correlation between CSF Aβ and atrophy in the precuneus. Our lack of statistically significant results in the medial temporal regions is not consistent with a body of evidence that CSF biomarkers are related to morphometric variability in the limbic regions, including the hippocampus [26, 27]. Our general finding of a dissociation between P-tau and other CSF biomarkers, however, is consistent with previous research showing preferential relationships between P-tau and brain atrophy assessed via MRI [28-30].

The preferential associations between biomarkers and structurofunctional change in the precuneus are consistent with the known distribution of AD pathology. The precuneus is the region of cortex occupying the medial aspect of the superior parietal lobule and lies adjacent to the posterior cingulate cortex [31]. Together, the precuneus and posterior cingulate regions are sites of early Aβ accumulation [32]. In contrast, the formation of tau-related pathology in this region occurs later in the pathogenesis of the disease [1, 33]. A number of in vivo imaging studies have confirmed the precuneus and posterior cingulate region as having affinity for Aβ ligands in patients with DAT and MCI [34, 35]. Imaging investigations in patients with DAT have revealed hypometabolism, hypoperfusion, structural atrophy, and abnormal functional connectivity in this combined region [36].

There is evidence to suggest two possible mechanisms of structurofunctional change in the precuneus and posterior cingulate region. Specifically, such changes may relate to local Aβ accumulation, or they may occur secondarily to disconnection from the mesial temporal region. The first mechanism, that changes in the precuneus and posterior cingulate regions may be secondary to local Aβ deposition, is consistent with the direct pathway of Aβ toxicity implicated in the amyloid cascade hypothesis [37]. A number of studies have reported that Aβ oligomers directly disrupt synaptic function, supporting the view that local Aβ deposition in cortical regions may result in local synaptic disruption, eventually leading to structurofunctional changes [38, 39]. A recent imaging study has confirmed the relationship between local Aβ deposition and structural atrophy, but only in regions of greatest Aβ deposition, including the precuneus and posterior cingulate [40]. There is also evidence that Aβ deposition precedes hypometabolism in the precuneus and posterior cingulate region, further supporting a relationship between local Aβ deposition and local structurofunctional change [41].

Secondary disconnection from mesial temporal regions is the second possible mechanism of structurofunctional change in the precuneus and posterior cingulate region. This hypothesis is supported by reports that the relative degree of hypometabolism exceeds the degree of structural atrophy in the precuneus and posterior cingulate region [42]. An implication of this discrepancy is that local structural atrophy in this region may be insufficient to explain the degree of hypometabolism, suggesting an additional contribution. A candidate for this additional contribution is that hippocampal atrophy induces a progressive breakdown of cingulum fibres, which then leads to precuneus and posterior cingulate hypometabolism [43]. In this sense, Wallerian degeneration might be an additional cause of hypometabolism and structural atrophy observed in this region [44].

It is still unclear as to whether the two mechanisms described above are equally contributory, or whether one or the other predominates at different stages of the disease. While the present finding of a relationship between Aβ-related (but not P-tau related) pathology and structurofunctional changes in the precuneus does not conclusively resolve this issue, it does strengthen the view that local Aβ deposition plays a significant role.

In terms of cognition, the precuneus is involved in a range of integrative cognitive functions in healthy adults, including episodic memory retrieval, self-processing, and visuospatial imagery, amongst others [31]. There is a body of evidence suggesting this region functions as the main `hub’ in the default mode network (DMN), a network of intrinsic functional connectivity that becomes prominently synchronised during rest [45-47]. Intrinsic functional connectivity in the DMN is abnormal in AD [48] and there is evidence to suggest this may be related to the degree of Aβ pathology [49]. Despite growing interest, the precise relationship between disruption to this region and the neurocognitive phenotype of Aβ is yet to emerge [50].

The finding of similar differential associations between Aβ and P-tau in the lateral temporal regions is also consistent with the proposed model. Like the precuneus, this region is the site of early Aβ deposition, whereas tau-related pathology does not become prominent until relatively later in the disease [1, 32]. Aβ deposition in this region has been well documented using in vivo) imaging approaches [35], and global Aβ affinity has been associated with structural atrophy in temporal neocortex [51]. In terms of the cognitive phenotype of DAT, hypometabolism and structural atrophy in this region has been associated with semantico-linguistic deficits [52]. Critically for the present study, local Aβ deposition in the superior temporal region has been specifically associated with the early disruption of language networks in DAT [21].

It was expected that, while Aβ-related pathology would be most strongly associated with markers of pathology in neocortical regions, structuro-functional changes in the mesial temporal regions would be most strongly related to levels of P-tau. The selective association between Aβ and neocortex was consistent with the first prediction, however the mesial temporal hypotheses were not supported. A reason for this might be related to the technical challenges of imaging mesial temporal structures. Automated parcellation and segmentation offer efficient and reliable means of examining volumes and metabolic activity of specific brain structures. There is some evidence that manual segmentation might be optimal for delineating mesial temporal structures, especially in the context of structural atrophy [53]. Manual segmentation, however, is time-consuming, and there are as yet no universally accepted protocols for hippocampal tracing [54].

Given that our sample is relatively young in terms of AD populations, it is possible that the development of tau-related pathology was not as advanced as the development of amyloid-related pathology. This might further explain why relationships were observed for amyloid, but not tau. A clear direction for future research is to investigate this hypothesis across a range of ages and disease stages. Further, given the sample size we were unable to explicitly model the effects of relevant comorbidities (such as vascular burden, cognitive function, and treatment status). As such, these findings should be viewed as hypothesis generating and future research should confirm our findings in larger cohorts which enable statistical adjustment of these potential confounds.

Only the relationship between amyloid-pathology and hypometabolism in the precuneus survived when only participants with ‘abnormal’ CSF Aβ concentrations were included in the analysis. The smaller number of participants in this sub-analysis (only 29 of the total sample of 39) raises the question of inadequate statistical power. Future, well-powered studies are required to fully confirm these findings while taking diagnosis CSF status into account. It is important to note, however, that the development of AD biomarkers is an insidious process that might begin decades before a clinical diagnosis is warranted [55]. For example, Braak and Del Tredici [56] have documented the accumulation of AD pathology as early as the first decades of life. In previous work, we have shown differential relationships between CSF biomarkers and cognition in patients with mild cognitive impairment [7]. As such, it might be more ecologically valid to consider the effects of CSF biomarkers across a range of clinical stages.

Hypometabolism in the mesial temporal regions is often difficult to identify [57]. One reason for this is the partial volume effect (PVE), which occurs when signal from a small structure is underestimated because voxels within it overlap more than one tissue class [58]. This effect occurs for structures smaller than two times the full-width half maximum (FWHM) of the scanner resolution, and is especially pronounced for GM structures adjacent to CSF spaces, such as the hippocampus [59]. While methods for correcting for PVE are available, they involve potential degradation of the data and can produce physiologically implausible results in the context of progressive atrophy [60].

The novelty of these findings rests on the direct investigation of the *differential* relationships between the cardinal pathologies and structure-functional changes. While previous work has reported different patterns of correlations between these variables, few have directly examined the differential nature of the relationships. In this study, we built upon our previous work developing a novel hypothesis of differential association between the cardinal pathologies and structural-functional changes [6, 7]. We then directly examined this hypothesis statistically by testing the *differences* between the relationships (i.e., by demonstrating a statistical interaction). We believe that this approach represents a substantial and novel contribution to the scientific literature.

Taken together, these findings support the proposed preferential association between pathological processes in AD and their topological distribution in the brain. The finding that hypometabolism and structural atrophy in the precuneus and lateral temporal regions were associated with Aβ but not P-tau supports the hypothesis that these two pathologies exert differential influence on neocortical regions. A limitation of this study was that it did not include healthy participants. An important task for future research will be to examine the relationships between biomarkers and cortical thickness measurements to ensure they are markers of pathological status, and are not otherwise related to brain structure and function. Cognitive markers were not evaluated as part of this study, as the primary focus was on the relationship between biomarkers and structuro-functional changes. This is an important limitation, and future work is necessary to examine the relationship between preferential biomarker changes and cognition. Although cognition was not directly evaluated in this study, these findings lend support to the hypothesis of dissociation between the two pathological processes in AD and their effects on the neurocognitive phenotype.

## Acknowledgements

Charles B Malpas was supported by an Alzheimer’s Australia Research Foundation Viertel postgraduate research scholarship during his doctoral candidature. He is currently supported by a National Health and Medical Research Council (NHMRC) Peter Doherty Australian Biomedical Fellowship.

